# Plasticity of Mitochondrial DNA Inheritance and its Impact on Nuclear Gene Transcription in Yeast Hybrids

**DOI:** 10.1101/394858

**Authors:** Sarah K. Hewitt, Kobchai Duangrattanalert, Tim Burgis, Leo A.H. Zeef, Daniela Delneri

**Author notes:** Corresponding author: (DD). These authors contributed equally to this work (Sarah K. Hewitt, Kobchai Duangrattanalert).

## Abstract

Mitochondrial DNA (mtDNA) in budding yeast is biparentally inherited, but colonies rapidly lose one type of parental mtDNA, becoming homoplasmic. Therefore, hybrids between different yeast species possess two homologous nuclear genomes, but only one type of mitochondrial DNA. We hypothesise that the choice of mtDNA retention is influenced by its contribution to hybrid fitness in different environments, and that the allelic expression of the two nuclear sub-genomes is affected by the presence of different mtDNAs in hybrids. Here, we crossed *Saccharomyces cerevisiae* with *S. uvarum* under different environmental conditions and examined the plasticity of the retention of mtDNA in each hybrid. We showed that on fermentable carbon sources at warm temperatures each parental mtDNA was equally likely to be retained, while at colder temperatures, hybrids preferentially retained mtDNA derived from *S. uvarum*. On a non-fermentable carbon source, hybrids retained *S. cerevisiae* mtDNA, independent of temperature. By acquiring transcriptome data and co-expression profiles for hybrids harbouring different mtDNA in a selection of environments, we found a clear pattern of concerted allelic transcription of one or the other sub-genome for specific biological pathways, supporting the notion that the hybrid cell works preferentially with one set of parental alleles or the other according to specific cellular functions. We argue that the type of mtDNA retained in hybrids affects the expression of the nuclear genome and the organism fitness in different environments, and therefore may have a role in driving the evolution of the hybrid nuclear genome in terms of gene retention and loss.

## Introduction

Mitochondria are dynamic and integral components of the cell, harbouring their own small genome of eight genes, but deriving the remaining ∼700 associated proteins from the nuclear genome (Prokisch et al. 2004). Mitochondrial DNA (mtDNA) is inherited in a non-mendelian manner outside of the nucleus. Hybrids between *Saccharomycetes* tend to carry mtDNA from just one parent *i.e.* they are homoplasmic. A homoplasmic state is common among many other eukaryotes, including humans, who inherit mitochondria from the mother’s gamete (Giles et al. 1980). When yeast cells mate, they fuse both their cell membrane, nuclear membrane, and the mitochondrial membranes from each parent (Nunnari et al. 1997; Berger and Yaffe 2000). The initial zygote harbours mtDNA from both parents, but over the course of few generations, mtDNA from one parent is lost whilst the other is retained (Marinoni et al. 1999; Verspohl et al. 2018).

All strains of *Saccharomyces pastorianus*, a natural lager brewing yeast hybrid between *Saccharomyces cerevisiae* and *Saccharomyces eubayanus* (Libkind et al. 2011), contain only mtDNA from the cryotolerant parent, *S. eubayanus* (Dunn & Sherlock 2008; Ranieri et al. 2008; Peris et al. 2014). Other industrial *Saccharomyces* brewery hybrids also retain the non-*S. cerevisiae* mitochondrial genome (Masneuf et al. 1998; Gonzáles et al 2006). The influence of environmental conditions on the inheritance of mtDNA in yeast and the impact of mtDNA on the fitness and transcription of yeast hybrids living and reproducing in different environments has been largely overlooked. Adaptation to temperature is one of the primary drivers of evolution in yeast (Salvadó et al. 2011) and is responsible for much of the phenotypic divergence between yeasts, even allowing different strains to exist in the same ecological niche (Sampaio et al. 2008). Thus, the environmental temperature present at the time of hybrid formation may play a role in mitochondrial genome choice, and this choice subsequently may affect the fitness of the organism. *Saccharomyces* yeast can both respire, which requires functioning mitochondria, or ferment, which does not. In fact, yeast can survive adequately without functioning mitochondria, and form smaller “petite” colonies. Thus, the type of carbon source present in the environment; *i.e.* a fermentable carbon source or a carbon source that can only be respired, may also affect mtDNA choice and the future fitness of the organism.

To better understand how environmental conditions may affect the inheritance of mitochondrial DNA and its impact on fitness and evolution of the hybrid genome (Hewitt et al, 2015), we crossed haploid *Saccharomyces cerevisiae* BY4741 (Sc) with spores of the cryotolerant species *Saccharomyces uvarum* NCYC 2669 (Su) under a range of temperatures and under either fermentation or respiratory conditions.

Given the fact that most mitochondrial proteins are nuclearly-encoded, we analysed the genome expression in different hybrid mitotypes and showed that the type of mitochondrial DNA present may create a nuclear allelic expression bias in different biological pathways that depends also on environmental conditions. Our expression analysis also reveals how different mtDNA may be affecting the hybrid’s transcriptome and how these expression differences may partially explain the differences in fitness observed between hybrids harbouring different mtDNA. Furthermore, comparing *S. cerevisiae* and *S. uvarum* alleles within the same hybrid background, and comparing alleles in hybrids compared to their parents allowed us to distinguish cis-regulatory and trans-regulatory differences, respectively.

## Results

### Construction of *Saccharomyces* hybrids under different environmental conditions and analysis of mitochondrial DNA retention

We investigated the ability of *S. cerevisiae* and the cryotolerant *S. uvarum* to successfully form viable hybrids (hereafter termed Sc/Su hybrids), under different environmental conditions. Using a micromanipulator, crosses were made on yeast-peptone agar plates containing a carbon source of either glucose (YPD, fermentable) or glycerol (YP + glycerol, non-fermentable) at 28 °C, 16 °C or 10 °C. The presence of both *S. cerevisiae* and *S. uvarum* subgenomes in the hybrids was confirmed using species-specific primers (Supplementary Fig S1). Additionally, the ploidy of all hybrids used in subsequent growth assays was confirmed by Sytox Green staining of DNA and analysis by flow cytometry (Supplementary Fig S2) (Haase 2004).

The percentage of viable hybrids created from crosses on YPD at 28 °C was just below 100 %. With each decrease in temperature, 16 °C and 10 °C, there was a sharp decrease in the percentage of hybrids that were successfully formed (73 % and 35 % respectively, Fig 1). The percentage of viable crosses generated on YP + glycerol was lower than those generated on YPD. Like crosses made on fermentative media, with each decrease in temperature there was a decrease in the number of crosses that resulted in viable hybrids. At 28 °C and 16 °C, 86 % and 23 % of crosses generated hybrids respectively. At 10 °C on YP + glycerol, less than 2 % of crossed parental strains successfully formed viable hybrids.

**Fig 1.**
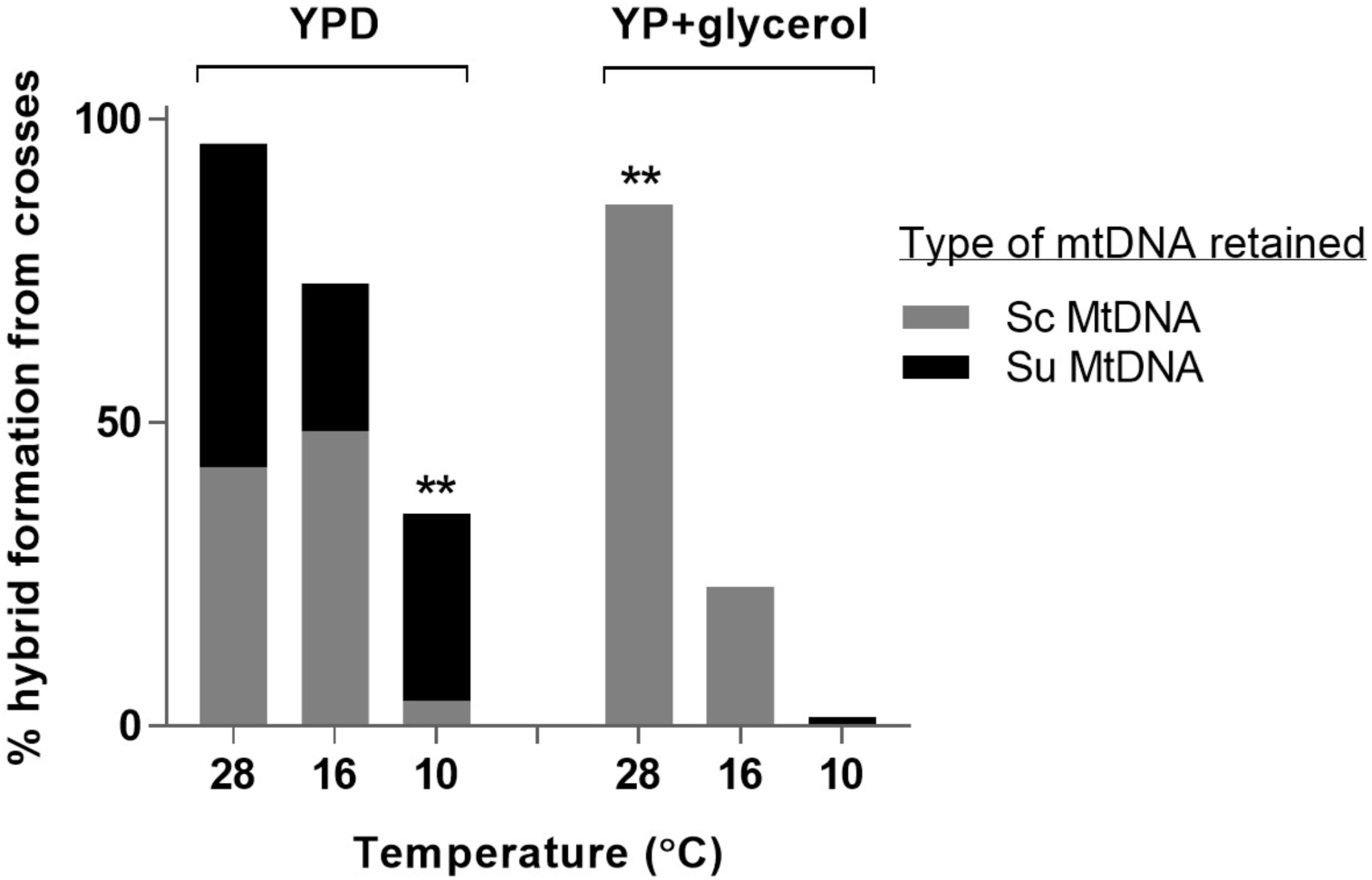
Percentage of hybrids that were successfully created from individual crosses under different laboratory conditions and the type of mtDNA they inherited. The number of expected hybrids was half the number of crosses made, given the random mix of MAT**a** and MATα *S. uvarum* spores. Sc mtDNA, mitochondrial genes show *S. cerevisiae* pattern; Su mtDNA, mitochondrial genes show *S. uvarum* pattern. ** p ≤ 0.01 within each carbon source (YPD, fermentable carbon source; YP, respiratory carbon source), exact binomial test of goodness-of-fit.

To determine which parental mtDNA was retained by each newly-constructed Sc/Su hybrid, two mitochondrial genes, *COX2* and *COX3*, were amplified by PCR and analysed by restriction length fragment polymorphism (RFLP). Furthermore, mitotypes were validated by Sanger sequencing of six mitochondrial genes *ATP6, COX1, COX2, COX3, COB*, and *SCE1* (Supplementary Fig S3). On the fermentable carbon source (YPD) at 28 °C and 16 °C, there was no significant difference between the number of hybrids that retained *S. cerevisiae* mtDNA and the number that retained *S. uvarum* mtDNA (Fig 1). However, on YPD agar at 10 °C the majority of hybrids retained *S. uvarum* mtDNA rather than *S. cerevisiae* mtDNA. Strikingly, on YP + glycerol, at 28 °C, all hybrids retained their mtDNA from *S. cerevisiae*, whilst all five hybrids created at 16 °C harboured *S. cerevisiae* mtDNA, while the single hybrid successfully formed at 10 °C contained *S. uvarum* mtDNA.

### The fitness of Sc/Su hybrid mitotypes varies with environmental condition

To determine the effect of mitotype on fitness, we assayed the growth of three hybrids with *S. cerevisiae* mtDNA (HMtSc) and three hybrids with *S. uvarum* mtDNA (HMtSu), under a variety of conditions. At 28 °C, the growth rate of HMtSc hybrids was faster than the growth rate of HMtSu hybrids in both carbon sources (Fig 2), although the magnitude of growth rate difference was greater under respiratory conditions (YP + glycerol). Strikingly, at 28 °C HMtSc grew faster than the parental strains, particularly in YP + glycerol. Growth rate difference between hybrids was most similar at 16 °C. Moreover, both hybrid mitotypes were fitter than the parents on both carbon sources at 16 °C (Fig 2). Interestingly, at 10 °C, HMtSu grew faster than HMtSc when inoculated on YP + glycerol, but not YPD. Finally, on YP + glycerol at 4 °C the only strains able to successfully grow were HMtSu and the *S. uvarum* parent, whilst on YPD, these strains showed superior growth to the *S. cerevisiae* parent and HMtSc. Overall, at warm temperature, HMtSc was fitter than HMtSu on both a respiratory and fermentable carbon source, whilst at cold temperature HMtSu was fitter than HMtSc on both a respiratory and fermentable carbon source. Fitness differences between strains were particularly pronounced on respiratory conditions.

**Fig 2.**
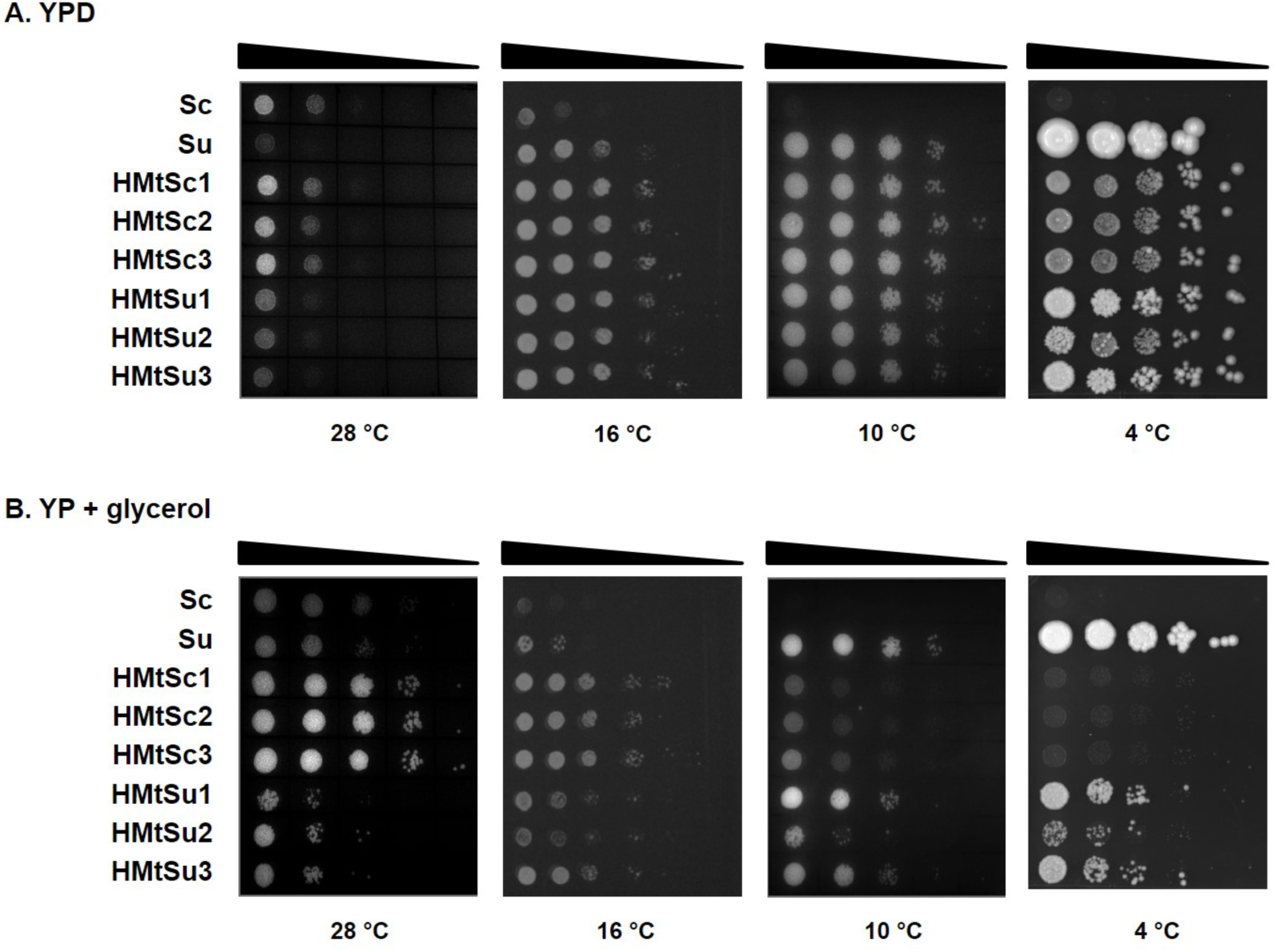
Growth assays of hybrid mitotypes on fermentative (YPD) or respiratory (YP + glycerol) carbon sources. The growth of three biological replicates of each hybrid mitotype (hybrids with *S. cerevisiae* mitochondria (HMtSc) and hybrids with *S. uvarum* mitochondria (HMtSu)) and the parental strains *S. cerevisiae* BY4743 (Sc) and *S. uvarum* NCYC 2669 (*Su*) on YPD (A) or YP + glycerol (B), at 28 °C, 16 °C, 10 °C or 4 °C and imaged every 12 hours. Each sample was diluted 1/10 up to a dilution of 1/10,000, moving from left to right, from an initial OD_600_ of 0.4 (far left). Images show time point of greatest growth differentiation between strains.

### RNA sequencing strategy and hierarchy of factors affecting transcriptional responses

Yeast mitochondria are comprised of proteins derived from both mtDNA (8 protein coding genes) and the nuclear genome ca. 700 genes (Prokisch et al. 2004). It is largely unclear how the mitochondrial DNA present in yeast hybrids influences the transcriptional response of the nuclear genome. We chose six independently-constructed hybrids of *S. cerevisiae* and *S. uvarum*, three with *S. cerevisiae* mtDNA and three with *S. uvarum* mtDNA for transcriptome analysis. These hybrids, along with the parental strains, were grown in either YPD or YP + glycerol at 28 °C, a temperature at which there was a significant phenotypic difference between mitotypes, and at 16 °C where both mitotypes showed a more similar fitness profile.

A strategy was designed to perform gene expression analysis of each subgenome in the hybrid strains. Firstly, reads were mapped to a combined genome index of *S. cerevisiae* (SacCer3) and *S. uvarum* (adapted from Scannell et al. 2011), only counting reads that mapped perfectly once to the combined genomes (Fig 3). Reads that were not uniquely mapped were discarded because it could not be established which subgenome they had originated from. The same workflow was applied to the parental strains to ensure that differential expression between all samples was not biased by excluding ambiguous reads. Secondly, reads were only counted for 5229 genes that had a clear one-to-one orthologous relationship between the two parental strains.

**Fig 3.**
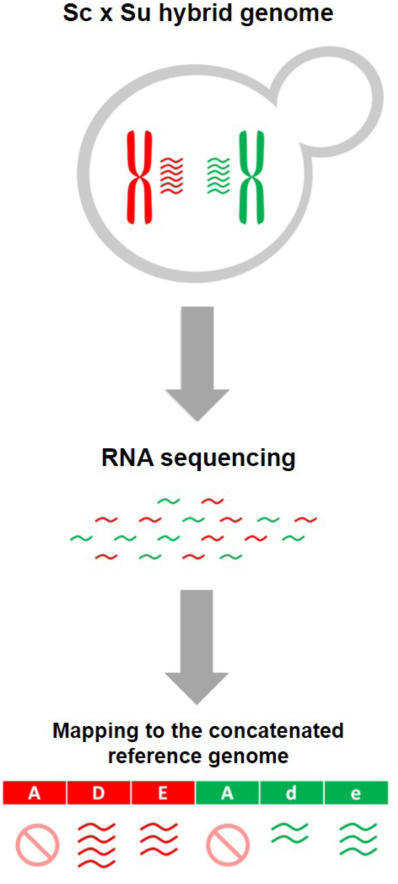
Strategy of RNA-seq data analysis to allow gene expression analysis of each subgenome in the hybrid strains. Reads were mapped to a combined genome index of *S. cerevisiae* (red colour) and *S. uvarum* (green colour) only allowing reads that mapped perfectly once to the combined genomes (i.e. D, E, d, e). Reads that were not uniquely mapped (i.e. A) were discarded.

Bioinformatic integration of gene expression from these two subgenomes was successful, based on a PCA analysis of the whole dataset. These data show that the biological factor of media type was the strongest contributor to gene expression, separating on the first component, whilst the *S. cerevisiae* and *S. uvarum* subgenomes separate on the second principal component (Fig 4A). To visualize the hierarchy of effects of the four experimental factors (medium, genome, temperature, strain), gene expression data were clustered (Fig 4B). Generally, within each subgenome, temperature has a greater impact on gene expression than the genetic background (hybrid-or parent-derived subgenome). The subgenomes of hybrids with different mtDNA cluster closer together than they do with either parent, with two exceptions: *i.* the *S. cerevisiae* subgenome of HMtSc in YP + glycerol at 28 °C clusters with its Sc parent; and *ii.* the *S. uvarum* subgenome of HMtSc in YPD at 28 °C clusters with its Su parent. This indicates that, in general, medium, subgenome type, and temperature have a greater influence on gene expression than the type of mtDNA in a hybrid, and the expression of a given subgenome will be more similar between mitotypes than between hybrids and parent.

**Fig 4.**
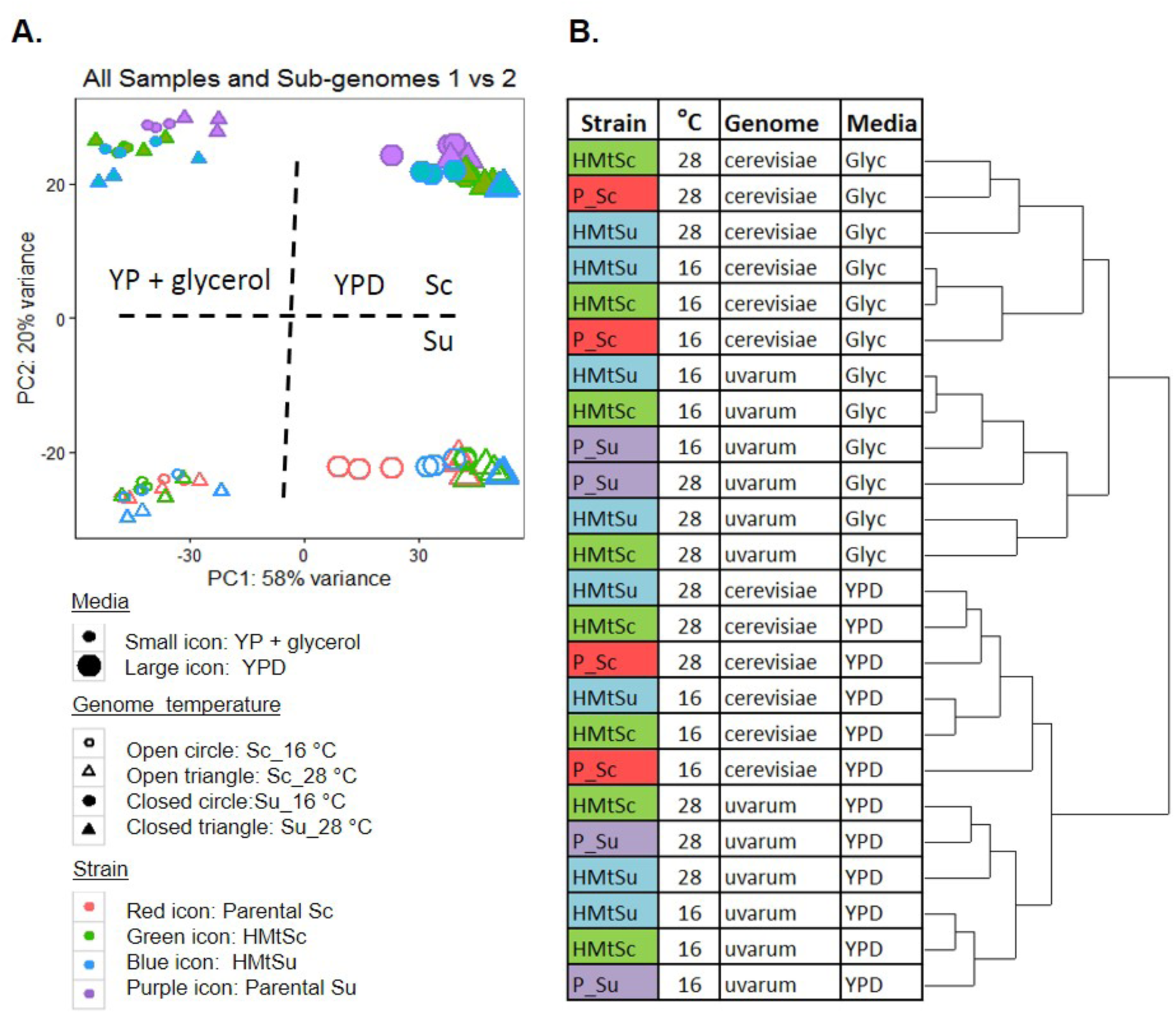
Clustering of expression data from RNAseq of hybrids and parental strains in different growth conditions. A. PCA analysis of the transcriptome of each subgenome of HMtSc and HMtSu, and the parental strain BY4743 *S. cerevisiae* and *S. uvarum* strain NCYC 2669. The expression data separates primarily by growth media (YP + glycerol or YPD), and secondly by subgenome. Dotted lines show the approximate main separation of clusters along the first and second principal components. B. Dendrogram describing more detailed clustering of strain and temperature within each growth condition.

### Plasticity of allele specific expression in Sc/Su hybrids

We investigated how strongly the expression levels of the Sc subgenome and Su subgenome are affected by the hybrid background, in relation to the environment. Specifically, we compared the gene expression of *S. cerevisiae* or *S. uvarum* alleles in the hybrids with their original expression profile in the respective parental backgrounds, within four growth conditions (Fig 5 and Table S1). The proportion of genes that were differentially expressed between parental strains and each hybrid depended greatly on growth condition and the mtDNA background (mitotype) of the hybrid. For example, there was a greater level of differential expression between the parental *S. cerevisiae* and the Sc alleles in HMtSc in cold conditions than warm conditions, regardless of carbon source (Fig 5 and Table S1). These expressional changes at cold are however mitigated in the HMtSu background. On the other hand, a higher proportion of genes were differentially expressed between parental *S. uvarum* and the Su alleles in HMtSc in the warm respiratory condition compared to the cold condition, and these changes were exacerbated in the HMtSu background (Fig 5 and Table S1). Overall, these data show that in the hybrids more alleles are affected at 28 °C when the hybrids contain Su mitochondrial DNA.

**Fig 5.**
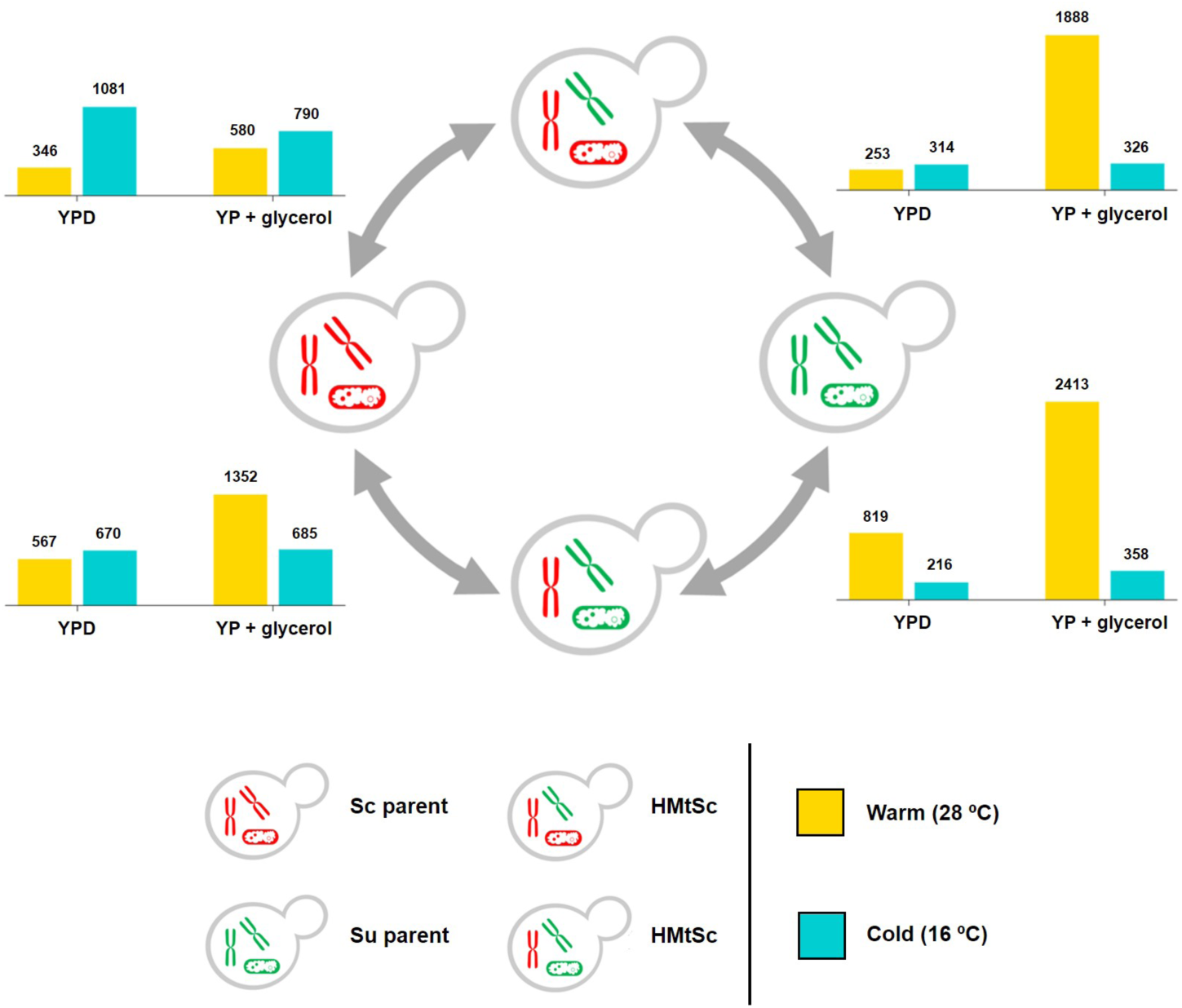
Differential expression between parental strains and hybrids. The number of differentially expressed genes between parental strains and the corresponding hybrid subgenome in HMtSc and HMtSu, grown in different conditions. Yellow bars denote growth conditions of 28 °C and blue bars denote growth conditions of 16 °C. YPD, fermentable carbon source; YP + glycerol, respiratory carbon source. Sc, *S. cerevisiae*; Su, *S. uvarum*.

We determined whether any groups of differentially expressed alleles were enriched for genes belonging to specific functional categories (Fig 6 and Table S2). Based on co-clustering statistics, we found that a large proportion of gene expression changes between hybrids and parents were common between the two mitotypes and therefore likely to be caused by the hybridisation event itself. Transcriptional shock upon hybridisation has been documented in various plants and animals, and is interestingly associated with differentiation of phenotypic traits in a wide range of environments (Hegarty et al. 2008; Wu et al. 2015; Wu et al. 2016; Ortiz-Merino et al. 2018). However, for those genes whose expression was significantly different in the two mitotypes, we performed an enrichment analysis to determine the key molecular pathways which are affected by the mitochondrial DNA (Table S2). The differential gene expression and preservation in the hybrids HMtSc and HMtSu was represented as circularised heat maps (Fig. 6). For the *Sc* subgenome (Fig. 6A) and Su subgenome (Fig 6B), we found four and three clusters, respectively, containing alleles differentially expressed between parents and hybrids that also differed significantly between mitotypes. Specifically, in the hybrid HMtSu in YP + glycerol at 28 °C, the Sc alleles within cluster 2 (Fig 6A) contained up-regulated genes in involved in cell wall remodelling (i.e. actin cortical patch and downstream signalling) and the Sc alleles within cluster 23 (Fig 6A) contained down-regulated genes involved in the cytoplasmic translation and mitochondrially-related functions (Table S2). It is known that the actin cytoskeleton plays an important role in several processes including endocytosis, cytokinesis, and cell morphology, and a role has been suggested in the regulation of cytoplasmic translation and mitochondrial function (Breitenbach et al. 2005; Boldogh and Pon 2006; Moseley and Goode 2006; Sattlegger et al. 2014). Indeed, these data are in accord with our phenotypic screening showing a poor growth of HMtSu on glycerol at 28 °C (Fig. 2).

**Fig 6.**
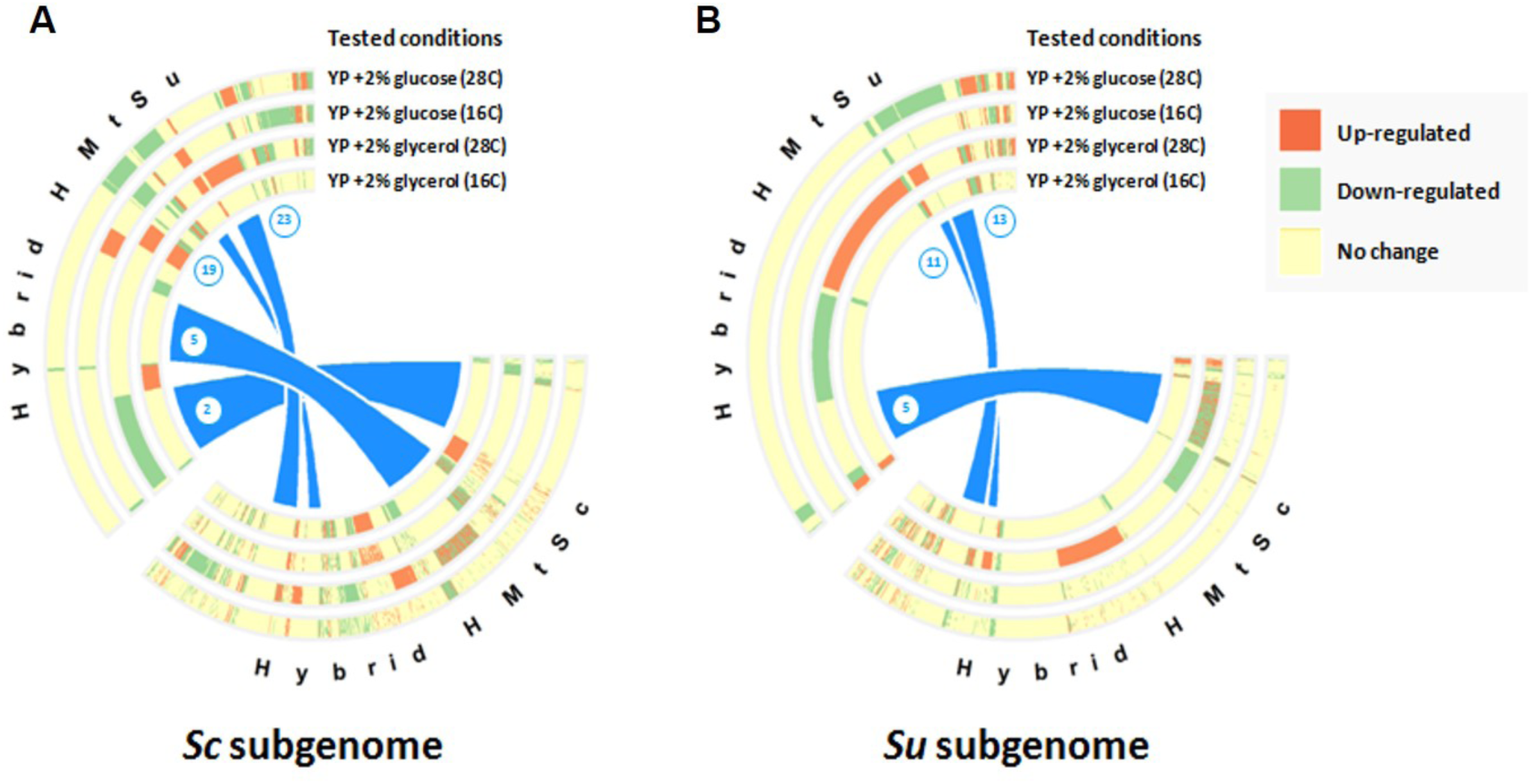
Circular heat maps representing differential gene expression and preservation of such changes in yeast hybrids. Circular heat map for the (A) *S. cerevisiae* (Sc) subgenome and (B) *S. uvarum* (Su) subgenome. Three different colours represent differential expression: red, up-regulation; green, down-regulation; light yellow, no significant change in gene expression. Heat maps of the two hybrids are connected to indicate divergence in differential expression of genes in two different mitotypes

In cluster 2, we also observed up-regulation of *RAS2, TPK2, PKH1, MSN2, SSK2*, and *SLT2* genes which are responsible for the propagation of stress signals along RAS and MAPK pathways (Chen and Thorner 2007; Tamanoi 2011). This suggests that the hybrid HMtSu is experiencing starvation, oxidative shock, and cell wall and membrane stresses when grown in glycerol at 28 °C. Despite cells trying to resolve the situation by promoting mitochondrial functions via actin and RAS signalling, Su mitochondrion appears less functional in the hybrid background when grown in warm conditions on a non-fermentable carbon source.

In YP + glycerol at 28 °C, the Su alleles within cluster 5 (Fig 6B) contains differentially regulated genes only in the HMtSc hybrid while in the HMtSu background, no alteration of expression was detected. The majority of these genes are involved in ribosome biogenesis and mitotic cell cycle control (Table S2).

### Differential expression of subgenomes in different hybrid mitotypes

We next compared patterns of gene expression directly between mitotypes HMtSc and HMtSu, by separately carrying out differential expression (DE) analysis on the Sc and Su subgenomes. Identification of any differences aimed to help clarify how harbouring different mtDNA may influence the expression of each subgenome within yeast hybrids. Almost half of the total alleles (Sc and Su alleles) have different expression between the two mitotypes in YP + glycerol at 28 °C (Table S3), indicating that the type of mtDNA harboured by the hybrid is extremely important for the nuclear transcription. These data agree with our phenotypic data: mitotypes were most separate in terms of growth at 28 °C, compared to 16 °C (Fig 2). In YPD at 28 °C, the type of mtDNA in the hybrid influenced the expression of the *S. uvarum* genome more than the expression of the *S. cerevisiae* subgenome. There were very few alleles significantly differentially expressed between mitotypes when hybrids were grown at cold, particularly respiratory conditions.

### Patterns of co-expression in HMtSc and HMtSu hybrids

Finally, we further examined the relationship between alleles by building a co-expression network and performing an unsupervised learning to cluster alleles with similar expression profiles across the four tested conditions (YP + 2 % glucose and YP + 2 % glycerol at 28 °C and 16 °C) and the two mitotypes (HMtSc and HMtSu). As shown in Fig 8, the network has around 1000 nodes and 3200 edges with its topology very close to scale-free. Interestingly, this topology is observed widely in biological systems such as metabolic and regulatory networks. To evaluate the clusters, we ran an exact binomial test and functional enrichment analysis to explain structural organisation by its allelic components and associated biological processes (Fig 8 and Table S4). The seven main clusters that showed similar expression levels are mainly made up of mitochondrial and ribosomal genes with different types of alleles distributed unevenly across different modules (Fig 8). Clusters with major functions in mitochondrial translation and endoplasmic reticulum membrane-related process are strongly enriched with *S. uvarum* alleles (more than 90 %; cluster 1 and cluster 2 in Fig 7) and clearly up-regulated in YP + glycerol medium at 28 °C with higher levels of expression in the HMtSu background. Genes involved in mitochondrial translation (cluster 1) were upregulated in YP + glycerol medium at 16 °C. These findings indicate the importance of these biological processes for metabolism of non-fermentable carbon source particularly at higher temperatures. Additionally, it is possible that such an increase in differential expression of the Su allele in these clusters may help explain fitness differences observed on solid media under the same conditions.

**Fig 7.**
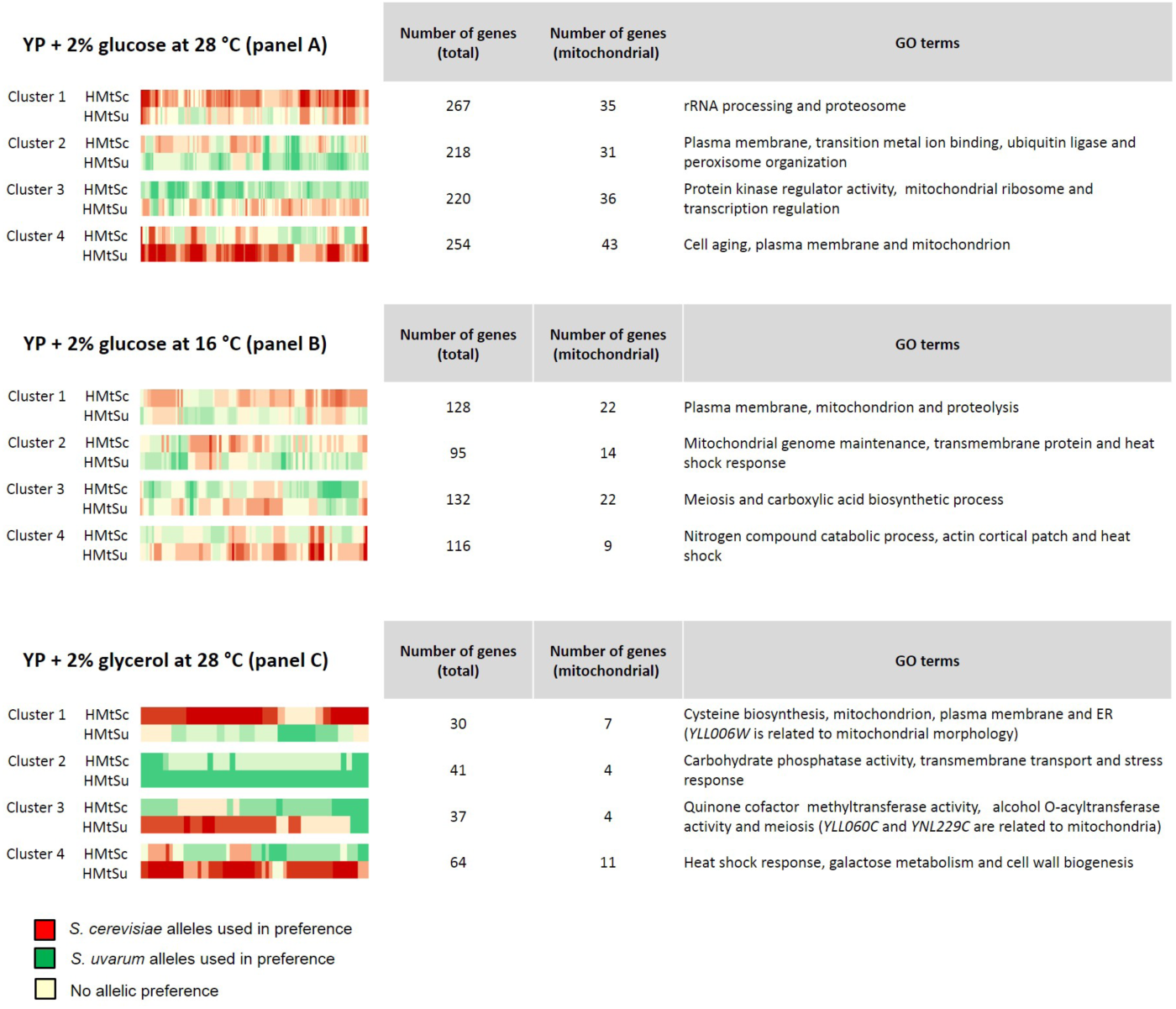
Clusters of genes in each condition where the locus expression ratio (Su vs Sc) changes between hybrids (HMtSc vs HMtSu). Each cluster represents a pattern of gene expression within each condition: Cluster 1, Sc alleles are upregulated compared to Su alleles in HMtSc but not HMtSu; Cluster 2, Su alleles are upregulated compared to Sc in HMtSu but not HMtSc; Cluster 3, Su alleles are upregulated compared to Sc alleles in HMtSc but not HMtSu and Cluster 4, Sc alleles are upregulated compared to Su alleles in HMtSu but not HMtSc. Number of genes in each cluster is shown along with the number of which are mitochondrial-encoded genes in parentheses. Gene ontology terms listed are those with fold-change of ≥ 2 at p ≤ 0.05.

**Fig 8.**
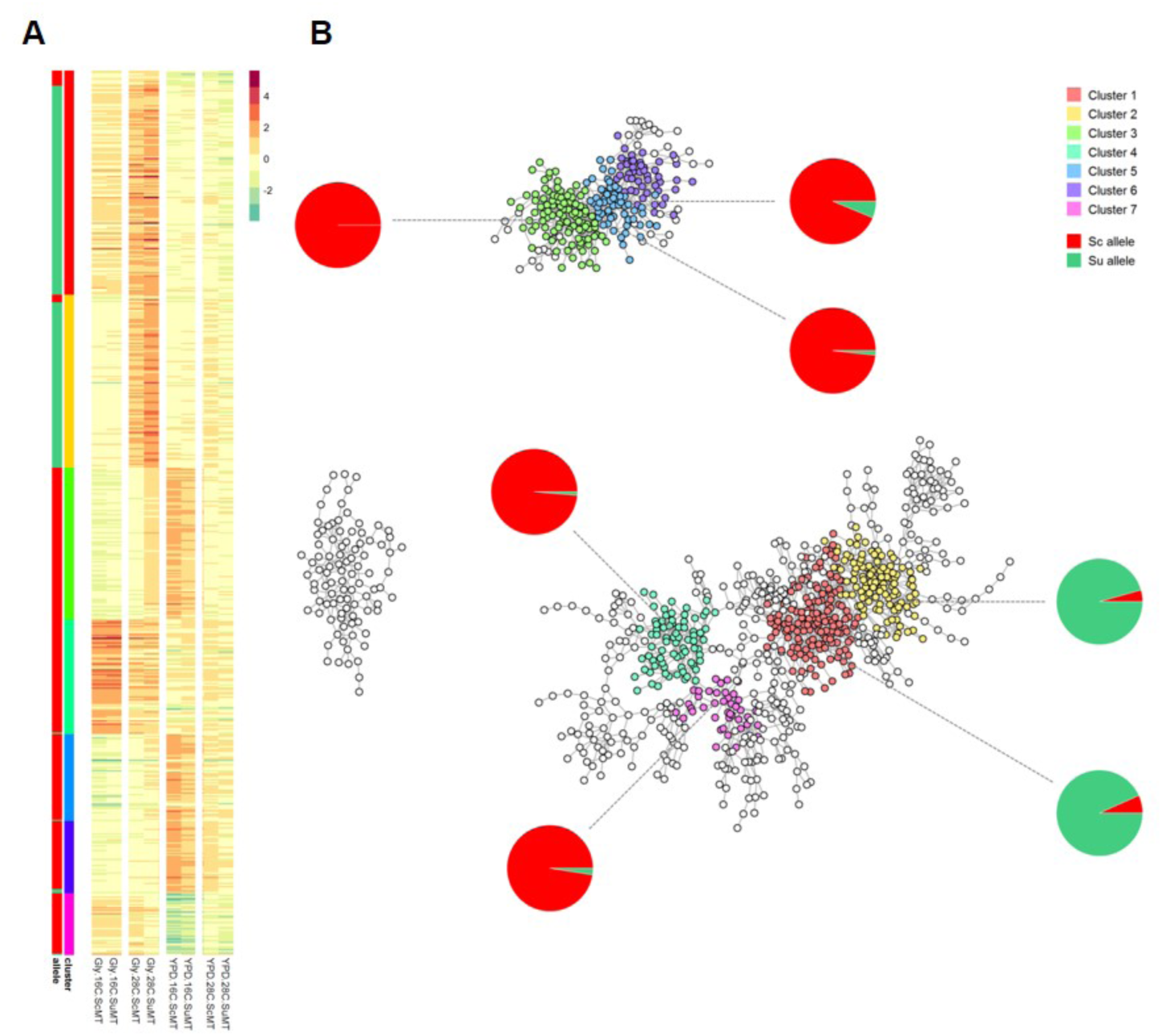
Analysis of allelic co-expression. (A) Heatmap was constructed based on significant correlation between alleles (i.e. Pearson coefficient ≥ 0.98 and p-value ≤ 0.05). Rows represent alleles of different genes and columns represent conditions used in this study. Each cell represents similarity in expression profiles with respect to the parental controls. Coloured bars show clusters identified by walktrap algorithm (step size = 13), which is implemented in the R package igraph, and proportion of Sc (red) and Su (green) alleles in the clusters. (B) A graphical representation of the co-expression map. The network was constructed based on significant correlation between alleles (i.e. Pearson coefficient ≥ 0.98 and p-value ≤ 0.05). Nodes represent alleles of different genes, and edges represent similarity in expression across the four testing conditions and the two hybrids. Clustering was performed using walktrap algorithm (step size =13) implemented in the R package igraph. Each cluster is complemented with a pie chart indicating the proportion of Sc (*S. cerevisiae*, red) and Su (*S. uvarum*, green) alleles.

In contrast to the first two clusters, we observed distinctively more Sc alleles in clusters 3, 5, and 6. These clusters are strongly associated with non-coding RNA processing and, interestingly, up-regulated in the HMtSc background at 16 °C. These data are partially expected, as similar events have been documented by several research groups. For example, it has been demonstrated that ribosome biogenesis is intrinsically cold-sensitive, as mutants with deleted genes related to this function are unable to grow efficiently at low temperatures (Moritz et al. 1991). Therefore, carrying Sc mitochondria may confer disadvantages to cold adaptation as certain genes were differentially regulated to deal with stress imposed by both mitochondria and temperature. More importantly, these results are in agreement with our phenotypic data, as the way yeast hybrid fitness is affected by cold conditions is akin to the growth defect in mutants with ribosome biogenesis malfunction. Cluster 4, encompassing mitochondrial translation and mitochondrial inner membrane-associated genes, has a great number of Sc alleles that were strongly upregulated in YP + glycerol at 16 °C, and in YPD at 16 °C in both hybrids. Cluster 7 was enriched for general RNA polymerase II transcription-associated Sc alleles and was largely downregulated in YPD at 16 °C in both hybrids.

## Discussion

### Hybridisation efficiency and mtDNA inheritance is influenced by carbon source and temperature

When yeast mate, mitochondrial proteins from each cell freely mix in the zygote, but their mtDNA remains largely segregated within mitochondrial membrane-bound nucleoprotein complexes called nucleoids (Berger and Yaffe 2000, Miyakawa et al. 1984; Azpiroz and Butow 1993; Kucej and Butow 2007; Solieri 2010). When the zygote later buds, the transmission of mtDNA depends largely on the position of the bud: an isogenic population will emerge from the distal end of the zygote, but mixed mtDNA may be present if the bud emerges from the medial portion (the point of mate fusion) of the zygote (Strausberg and Perlman 1978; Zinn et al. 1987). Thus, the transmission of mtDNA into individual buds deriving from a hybrid zygote is inherently non-random, due to the immobility of nucleoids (Nunnari et al. 1997; Okamoto et al. 1998). Despite the potential transmission of either or both mitochondrial genomes into new yeast buds, within 20 cell divisions populations of hybrid yeast deriving from a single mating become homoplasmic (Berger and Yaffe 2000;,Sherman, 1982). There is some evidence in *S. cerevisiae* that mitochondria of the highest quality, e.g. those that exhibit less damage by reactive oxygen species, are preferentially transmitted into new buds (Aguilaniu et al. 2003; Erjavec and Nyström 2007; Klinger et al. 2010). Our work investigated the effect of environmental conditions on the role of mtDNA inheritance and in the fitness of yeast hybrid mitotypes. We hypothesised that the type of mtDNA established in a population derived from a single hybrid cross may be a product of its fitness in particular environment, and may in turn affect the organisms’ fitness in future environments.

We crossed *S. cerevisiae* and *S. uvarum* haploids and cultured them under different environmental temperatures, supplying them with either a fermentative or respiratory carbon source. Hybrid yeast were more readily formed from crosses made in warmer fermentation conditions than those made in cooler respiratory conditions. Of those successfully formed hybrids, all hybrids grown with a respiratory carbon source, YP + glycerol, at 28 °C and 16 °C, inherited mtDNA from *S. cerevisiae*, whereas those constructed on YPD inherited mtDNA from either parent at 28 °C and 16 °C. Given that glucose in YPD is predominantly fermented, whereas glycerol in YP + glycerol is exclusively respired via the mitochondria, our results would therefore indicate that *S. cerevisiae* mtDNA is preferentially inherited over *S. uvarum* mtDNA where respiratory ability is crucial, at least under these conditions. These findings conform to the hypothesis by Albertin and coworkers (Albertin et al. 2013), who predicted using *in silico* modelling that a 1:1 mixed population of *S. cerevisiae* and *S. uvarum* mitotypes in respiratory conditions would, after several generations, be dominated by the *S. cerevisiae* mitotype. Studies on over 20 hybrids found that parental mtDNA types could be randomly inherited by *S. cerevisiae* × *S. uvarum* hybrids (Albertin et al. 2013; Solieri et al. 2008). Other smaller studies have also reported random inheritance of either parental mtDNA (Verspohl et al. 2018). However, a large scale project by Marinoni and coworkers (Marinoni et al. 1999) involving crosses from two different strain pairs observed a 100 % transmission rate of *S. cerevisiae* mitochondrial DNA under similar conditions. In another study, 28 hybrids were constructed and all harboured mtDNA derived from *S. uvarum* (Lee et al. 2008). The above-mentioned studies involved the production of yeast hybrids by employing standard *S. cerevisiae* laboratory conditions (*i.e.* in warm glucose-based media) to construct crosses, rather than a respiratory media, on which we almost exclusively observed *S. cerevisiae* mtDNA inheritance. There may be a strong strain dependence of mtDNA inheritance, a phenomenon also suggested by other groups (Solieri et al. 2008; Albertin et al. 2013) and supported by a recent study of mtDNA inheritance in crosses of *S. cerevisiae* and *Saccharomyces paradoxus* where both YPD and YP + glycerol were used as carbon source (Hsu and Chou 2017). Here they found that crossing one pair of *S. cerevisiae* and *S. paradoxus* strains in YPD (with 2 % glucose, as with our study) and YP + glycerol resulted in the retention of mtDNA from the *S. cerevisiae* parent, whilst crossing an alternative pair (different strains) of *S. cerevisiae* and *S. paradoxus*) resulted in the retention of mtDNA from the *S. paradoxus* parent, regardless of carbon source. However, in one pair of strains, increasing the percentage of glucose in YPD reversed preference from the *S. paradoxus* parent to the *S. cerevisiae* parent.

In our study, we additionally constructed hybrids at a variety of environmental temperatures. A previous study of *Cryptococcus neoformans* showed that temperature influenced the inheritance of mtDNA from the MATα or MATa parents, whereby lowering the temperature allowed increasingly fewer hybrids to inherit mtDNA from the MATα parent (Yan et al. 2007). In our study, at a cooler temperature of 10 °C we found that *S. uvarum* mtDNA was strongly preferentially retained in hybrids constructed on YPD agar, whilst the single successfully formed hybrid constructed at 10 °C in YP + glycerol also carried the same mtDNA. These data may provide the first clue as to why *S. eubayanus* (which, like *S. uvarum*, is a cold-favouring species), rather than *S. cerevisiae* mtDNA, is inherited in strains of the lager-brewing yeast *S. pastorianus* (Libkind et al. 2011; Dunn and Sherlock 2008; Ranieri et al. 2008).

### mtDNA influences the fitness of yeast hybrids in an environment-dependent manner

In an attempt to explain the mtDNA inheritance patterns we observed, we investigated whether hybrids with *S. cerevisiae* mtDNA were fitter than those with *S. uvarum* mtDNA, and if environmental conditions affected the outcome. We found that hybrids that had inherited *S. cerevisiae* mtDNA grew at least marginally better than those with *S. uvarum* mtDNA on fermentation media (YPD) at all temperatures in which we were successfully able to form hybrids and examine mtDNA inheritance, failing to explain the non-biased retention of mtDNA at 28 °C and 16 °C and the preference for *S. uvarum* mtDNA at 10 °C. However, we further looked at fitness at 4 °C, where we found that hybrids with *S. uvarum* mtDNA grew better than those with *S. cerevisiae* mtDNA.

When hybrids were grown on a respiratory carbon source, the differences in mitotype phenotype were generally much clearer than those observed when hybrids were grown on a fermentable carbon source, and hybrids with *S. uvarum* mtDNA also grew better than those with *S. cerevisiae* mtDNA at 10 °C as well as 4 °C. Our mtDNA inheritance results for YP + glycerol growth are in agreement with and potentially explained by fitness differences between the two mitotypes. However, it has been shown that differences between hybrids grown in fermentation conditions is not necessarily reflected by the type of mtDNA in the cell (Albertin et al. 2013; Origone et al. 2018). Still, Picazo and coworkers found that Sc/Su hybrids that inherited *S. cerevisiae* mtDNA showed an increased tolerance to oxidative stress and dehydration (Picazo et al. 2015). Other studies have indicated a possible role for mitochondria in cell growth, sugar uptake and drug tolerance in fermentation conditions and during the fermentation process, such as generating different organic acid profiles (Motomura et al. 2012; O’Connor-Cox et al. 1996). Researchers also found that fitness differences between *S. cerevisiae* × *S. paradoxus* mitotypes explained the inheritance of mtDNA in some conditions and not others, but that effects also varied depending on the strains used (Hsu and Chou 2017). Regardless of the factors affecting inheritance, the retention of *S. cerevisiae* mtDNA is evidently advantageous to hybrids growing in warm temperatures and particularly so in respiratory conditions, where mitochondrial function is essential. Indeed, previous studies have shown that hybrids with *S. cerevisiae* mtDNA are more likely to respire and less likely to ferment than hybrids with *S. uvarum* mtDNA (Solieri et al. 2008).

Physical differences of mtDNA between *S. cerevisiae* and *S. uvarum* could feasibly explain fitness differences. For example, *S. cerevisiae* has eight origins of replication whilst *S. uvarum* has four (Masneuf et al. 1998). However, this would fail to explain major differences seen between *S. cerevisiae* and *S. paradoxus* and different *Saccharomyces* strains of the same species (Hsu and Chou 2017). Furthermore, it is unclear which mechanisms may be acting on mtDNA selection associated with temperature variation. MtDNA divergence has been shown to be associated with climate in humans, and this variation may be driven by natural selection acting on mutations in mtDNA (Balloux et al. 2009). Human populations living in colder environments were shown to have lower mitochondrial diversity, whilst mtDNA *ND3* and *ATP6* genes appeared to have a clear association with temperature. MtDNA variation has also been linked to the adaptation to altitude in humans, particularly the Tibetan people, and birds (Kang et al. 2013; Zhou et al. 2014; Li et al. 2016). Mechanisms behind these adaptations include selection pressures on *ND2, ND4*, and *ATP6* genes in galliform birds; mutations in particular mtDNA genes in complex I of the inner mitochondrial membrane and selection for low mtDNA copy number.

From an industrial perspective, we suggest that the non-*S. cerevisiae* mitochondrial DNA, such as *S. uvarum* and *S. eubayanus*, of hybrid industrial strains used for fermentation has been selected for by a preference for growth in low temperature (particularly hybrid lager yeast strains; Masneuf et al 1998; Gonzáles et al. 2006) and decreased respiratory capacity compared to the *S. cerevisiae* mitochondria, favouring fermentation. At 16 °C, both hybrid mitotypes grew faster than both parental strains, whilst hybridisation was advantageous to *S. cerevisiae* at 4 °C if HMtSu is retained, and advantageous to *S. uvarum* at 28 °C if HMtSc is retained. Temperature may play a wider role in the likelihood of hybridisation both occurring and being evolutionary advantageous.

### Alleles show a high degree of plasticity of expression between hybrids and parents, between mitotypes, and within hybrids

The *S. uvarum* subgenome was more differentially expressed between hybrids and parents at warm temperatures, whilst the *S. cerevisiae* subgenome was more differentially expressed at cool temperatures. Both *S. cerevisiae* and *S. uvarum* alleles showed a high degree of expression plasticity between parental cells and hybrids, implying that the expression in hybrids may be subject to different trans-acting effects from each parental genome(de Meaux et al 2006; Guo et al. 2008). By this explanation, the effects of trans-acting regulation depended heavily on the environmental condition: *S. uvarum* alleles exerted a greater effect on the expression of *S. cerevisiae* allele in cool conditions than in warm conditions, whilst *S. cerevisiae* alleles exerted a greater effect on the expression of *S. uvarum* alleles in warm conditions than in cool conditions. Additionally, or alternatively, gene expression differences between parental strains and hybrid subgenomes may be due to differences in gene dosage from either the *S. cerevisiae* or *S. uvarum* genome when comparing diploid parents to the equivalent haploid subgenome in hybrids.

Strikingly, when directly comparing the expression profile of mitotypes, there were far more differentially expressed genes in warm conditions than cool, particularly between the two *S. uvarum* subgenomes, and particularly in YP + glycerol. In agreement, the growth of the two mitotypes on agar differed most greatly at 28 °C compared to 16 °C and particularly in YP + glycerol. In YP + glycerol at 28 °C, a large proportion of mitochondrial-associated Sc alleles are upregulated compared to Su alleles in HMtSc, but not in HMtSu. Expression data could therefore begin to explain growth differences between yeast hybrids with different parental mtDNA.

Differential expression within hybrids can be used to infer allele cis-regulatory effects, given that alleles within a hybrid share the same trans-acting background (de Meaux et al 2006; Guo et al. 2008). In our study, allele preference is probably more important in respiratory conditions than fermentation conditions, given the larger number of genes differentially expressed in these conditions. In both mitotypes, there is also a generally a stronger allele preference at 28 °C than at 16 °C within any given media and strikingly, in YP + glycerol at 16 °C, over a third of genes with a preference for the *S. cerevisiae* allele in both hybrids are mitochondrial-associated. A study of allele preference in crosses of *Arabidopsis thaliana* and *A. lyrata* revealed a strong preference for the expression of *A. lyrata* alleles (He et al. 2012). Almost 90 % of genes that showed allele specific expression had a preference for *A. lyrata*, likely caused by genome specific differences in epigenetic silencing. We found that different environmental conditions promoted differences in allele preference between hybrid mitotypes, but less strongly between subgenomes. The identification of preferred parental mitochondrial-associated alleles is particularly relevant in our study, where mtDNA inheritance in hybrids greatly impacts fitness under different conditions.

### Patterns of co-expression in Sc/Su hybrids across different environmental conditions

Alleles clustered based on similar levels of transcription showed an uneven distribution of Sc or Su alleles across different transcription clusters. Two clusters enriched for genes performing mitochondrial translation and endoplasmic reticulum membrane-related processes were comprised almost entirely of *S. uvarum* alleles. We found that both clusters were upregulated in YP + glycerol at 28 °C, and particularly in HMtSu, indicating the importance of these processes for metabolising non-fermentable carbon sources at high temperature. It also helps explain the differential expression of the *S. uvarum* alleles and slower growth of HMtSu in growth assays on YP + glycerol at 28 °C. In contrast to the first two clusters, cluster 3, 5, and 6 contain more Sc alleles and have a strong association with non-coding RNA processing. Several genes in these clusters are involved in the biosynthesis of ribosomal and transfer RNA. Interestingly, these genes were strongly up-regulated in HMtSc grown in YPD (YP + 2 % glucose) at 16 °C. It is known that ribosome genesis is intrinsically cold-sensitive, as ribosome genesis deletion mutants are unable to grow efficiently at low temperatures (Moritz et al. 1991). Transcriptional regulation was retained nearly exclusively within the same allele types, which could potentially be explained by divergence of cis-regulatory elements and mitochondrial genome of the two related yeast species. Assume that this regulatory variation is essential for optimal growth within their specific environments, combining the two genomes may increase a repertoire of functional abilities towards a wider range of stresses.

The work presented in this study sheds light on the plasticity of yeast hybrids in response to a range of environmental conditions and provides evidence that the choice of mtDNA retained by newly-formed hybrids could be evolutionarily advantageous. Our findings will aid the development of industrially important hybrid strains and help elucidate the evolutionary mechanisms of hybrid vigour.

## Materials and Methods

### Yeast strains and media

*Saccharomyces cerevisiae* BY4741 and *Saccharomyces cerevisiae* BY4743 were obtained from Thermo Scientific, UK. A KanMX cassette was previously engineered into the neutral *AAD3* locus of *S. cerevisiae* BY4741 (*MAT****a*** *his3Δ0 leu2Δ0 met15Δ0 ura3Δ0*) to confer selectivity (Piatkowska et al. 2013). *Saccharomyces uvarum* NCYC 2669 was obtained from the National Collection of Yeast Cultures, UK. The triploid hybrid used as a control strain for ploidy analysis was constructed previously (this lab) between *S. cerevisiae* BY4741 and *S. uvarum* NCYC 2669. Yeast strains were maintained on YPDA (2 % (w/v) Bacto Peptone, 1 % (w/v) Bacto Yeast Extract, 2 % (w/v) glucose, and 2 % (w/v) agar). Hybrid growth was assessed on either YPDA or YP + glycerol (2 % (w/v) Bacto Peptone, 1 % (w/v) Bacto Yeast Extract, 2 % (w/v) glycerol, and 2 % (w/v) agar).

### Construction of hybrids

*S. uvarum* NCYC 2669 was sporulated by inoculating into pre-sporulation media (0.8 % (w/v) Bacto Yeast Extract, 0.3 % (w/v) Bacto Peptone, and 40 % glucose) overnight, pelleting the cells and spreading thinly on minimal sporulation plates (1 % (w/v) potassium acetate and 2 % agar) and then incubating for five days at 16 °C, or until tetrads were observed microscopically. Tetrads were collected and incubated in 40 µL 1.5 M sorbitol at 37 °C for 7 minutes with 0.2 mg/mL Zymolyase (MP Biomedicals, USA) to disrupt the integrity of asci and release haploid spores. Individual *S. uvarum* NCYC 2669 mixed mating type spores and *S. cerevisiae* BY4741 *MAT***a** haploids were placed into physical contact for mating using a Singer Instruments MSM micromanipulator on a surface of either YPD agar or YP + glycerol agar. Plates were incubated at 10 °C, 16 °C, or 28 °C until colonies of 2-3 mm were visible. Colonies were then replica plated onto SDA + G418 selective plates (0.17 % (w/v) YNB without additional amino acids, 0.5 % (w/v) ammonium sulphate, 2 % (w/v) glucose, 2 % (w/v) agar, and 300 µg/mL G418 geneticin). These conditions exclusively select for hybrids, which can both tolerate geneticin and grow in the absence of additional amino acids. Plates were re-incubated until hybrid colonies were visible. The colonies were then re-streaked onto SDA + G418 selective media before testing. The hybrid strains passed through roughly 50 generations. Strains were stored at −80 °C in 15 % (v/v) glycerol.

### Whole genome extraction

Colonies were inoculated in YPD and grown at 30 °C, 200 rpm for 20 hours. Genomic DNA was extracted from yeast cells using Wizard Genomic DNA Purification Kit (Promega, UK) according to the supplied protocol and rehydrated in H_2_O overnight at 4 °C. The presence of genomic DNA was assessed by gel electrophoresis on a 1 % (w/v) agarose gel in TAE (40 mM Tris base, 20 mM acetic acid, and 1 mM EDTA) buffer. DNA was quantified using a Nanodrop 1000 spectrophotometer (Thermo Scientific, USA).

### Confirmation of hybrid genotype by PCR

Species-specific primers were used to check for the presence of *S. cerevisiae* and *S. uvarum* subgenomes within each putative hybrid colony (Piatkowska et al, 2013). Each reaction consisted of 100 ng genomic DNA, 4 ng of each upstream and downstream species-specific primer, 2 µL 5× MyTaq Reaction Buffer, 0.1 µL MyTaq DNA Polymerase, and up to 10 µL of H_2_O. Cycling conditions were as follows: 95 °C for 5 minutes; 35 × (95 °C for 45 s, 55 °C for 45 s, and 72 °C for 1 minute); final extension 72 °C for 8 minutes.

### Ploidy analysis by flow cytometry

Cells were prepared for flow cytometry as previously described [15]. Briefly, overnight cultures of yeast cells were fixed using 95 % ethanol, treated with 2 mg/mL RNAse (Sigma, USA) for two hours at 37 °C and 5 mg/mL protease solution for 30 minutes at 37 °C, stained with 1 µM Sytox Green (Life Technologies, USA) and sonicated using a Diagenode Biorupter (Diagenode Corporation, Belgium) for 20 seconds on low power to disrupt clumps. Cells were analysed with 488 nm excitation on a Beckman Coulter CyAn^™^ ADP flow cytometry machine (Beckman Coulter, USA). Green fluorescence was collected at 523 nm. Data were analysed using Summit v4.3 software (Beckman Coulter).

### Analysis of mitochondrial DNA origin by PCR

Primers for the amplification of mitochondrial *COX2* sequence were previously available (Belloch et al. 2000). All the other primers for amplifying *ATP6, COX1, COX3, COB,* and *SCE1* were designed using Primer3 (Koressaar and Remm 2007) to amplify the sequence from both parental species. PCR products were subsequently analysed by RFLP. Mitochondrial gene reactions consisted of 400 ng genomic DNA, 10 ng of each primer, 10 µL 5× MyTaq Reaction Buffer, 0.5 µL MyTaq DNA Polymerase, and up to 50 µL of H_2_O. The *COX2* gene was amplified as follows: 95 °C for 5 minutes; 45 × (94 °C for 40 s, 45 °C for 35 s, 72 °C for 35 s); final extension 72 °C for 10 minutes. The *COX3* gene was amplified as follows: 95 °C for 5 minutes; 45 × (94 °C for 40 s, 55 °C for 35 s, 72 °C for 35 s); final extension 72 °C for 10 minutes. Mitochondrial gene PCR products were purified using the Qiagen PCR Purification Kit and quantified using a Nanodrop 1000 spectrophotometer (Thermo Scientific, USA). To assess the parental origin of the amplified mitochondrial genes, 1000 ng of each was digested with either 1 U *Hin*f I and *Hin*d III, (Roche Diagnostics, Switzerland) for *COX2, COX3*, respectively, along with 2.5 µL SuRE/cut Buffer H (10×) and up to 25 µL H_2_O for 1 hour at 37 °C. PCR products and restriction enzyme digest products were analysed along with parental controls by electrophoresis on a 1.5 % (w/v) agarose TAE gel to determine their parental origin.

### Growth assays

Colonies were inoculated in 5 mL YPD overnight, washed twice in distilled sterile H_2_O and resuspended in 5 mL H_2_O to an OD_600_ of 0.4. Petri dishes (150 × 15 mm) were used to make solid media plates of YPD or YP + glycerol. 1/10 dilutions of each culture were made up to a dilution of 1/10,000. 4 µL of each culture and their three dilutions was spotted on to the agar. Plates were incubated at 4 °C, 10 °C, 16 °C, or 28 °C. Photographs of each plate were recorded every 12 hours using a Bio-rad Geldoc XR transilluminator with the Bio-Rad Quantity One software (Bio-Rad, USA). Note that diploid parental *S. cerevisiae* BY4743 was chosen rather than the parental haploid BY4741, to provide a ploidy comparative to the hybrids.

### RNA extraction and sequencing

Total RNA was extracted from three biological replicates of *S. cerevisiae* diploid BY4743 and *S. uvarum* diploid NCYC 2669, and from one biological replicate each of three independently constructed hybrids of BY4741 × NCYC 2669 with *S. cerevisiae* type mtDNA and three independently constructed hybrids of BY4741 × NCYC 2669 with *S. uvarum* mitochondria. Cultures were inoculated in either 5 mL of YPD or YP + glycerol and grown overnight at the appropriate temperature, 200 rpm. Cultures were scaled up in 50 mL fresh media at the appropriate temperature, 200 rpm to obtain mid log phase of between OD_600_ 0.5 and OD_600_ 0.8, at which cells were actively and rapidly replicating. To isolate RNA, cells were pelleted, flash frozen using liquid nitrogen and ground to a fine powder and pre-chilled to −80 °C. To break open the cell walls, Trizol (Ambion, Life Technologies, USA) was dispensed drop-wise onto the frozen pellet, which was then left to defrost. The thawed mixture was left for 5 minutes to allow dissociation of the nucleoprotein complexes. The mixture was pelleted for 10 minutes at 12,000 × g and 200 µL chloroform was added to every 1 mL of clear supernatant. Mixture was inverted and left for 10 minutes at room temperature. Phases were separated by centrifugation at 12,000 × g for 5 minutes. The upper clear aqueous RNA phase was transferred to fresh tubes and the RNA was precipitated with 1:1 isopropanol left for 10 minutes at room temperature. The mixture was centrifuged at 12,000 × g for 10 minutes and the supernatant was removed. The RNA pellet was washed with 1 mL (DEPC)-treated 70 % ethanol, air dried, dissolved in 1:1 lithium chloride (Ambion, Life Technologies, USA): DEPC-H_2_O and incubated overnight at −20 °C to precipitate a refined RNA pellet. The pellet was washed twice with DEPC-treated 70 % ethanol as before and then redissolved in 50 µL DEPC-H_2_O. The integrity of the RNA was checked on 1.5 % agarose gel. The quantity and quality of RNA was assessed using a Nanodrop 1000 spectrophotometer (Thermo Scientific, USA).

mRNA was sequenced on an Illumina Hiseq 2500 (Illumina, USA) platform. Library preparation and RNA-Seq was performed by the Genomic Technologies Core Facility at The University of Manchester, UK. A total of twelve samples were sequenced: three *S. cerevisiae* and three *S. uvarum* biological replicates; three independently constructed hybrids with *S. cerevisiae* mitotype (HMtSc); and three independently constructed hybrids with S. *uvarum* mitotype (HMtSu) for each temperature tested.

### Assembly, annotation, and DE analysis

*S. cerevisiae* (UCSC SacCer3) and *S. uvarum* genomes (Scannell et al. 2011) were used to create an artificial genome composed of both species. Sequenced single-end reads were mapped to this artificial genome using STAR mapper, reporting only reads that could be uniquely mapped to one place in the combined genomes. The number of reads mapped per *S. cerevisiae* and *S. uvarum* genes were counted with htseq. Counts were normalised to adjust for the total number of reads. DESeq (Anders et al. 2010) was used normalise data and to calculate differential expression and mean values for each strain.

Orthologous relationships between the 6010 annotated *S. bayanus var. uvarum* (now known as *S. uvarum*) genes (Scannell et al. 2011) and *S. cerevisiae* genes (SGD, www.yeastgenome.org) were calculated using Inparanoid (Remm et al. 2001). 5224 genes had a simple one-to-one orthologous relationship with a *S. cerevisiae* gene. A further 105 genes with a one-to-many relationship were assigned to best matched *S. cerevisiae* gene. These 5329 pairs of orthologs are the basis for the analyses presented here. The published assembly of *S. S. bayanus var. uvarum* (Scannell et al. 2011) appears to contain a section of the mitochondrial genome misassembled into nuclear contig Sbay_6. This erroneous section of Sbay_6 has been replaced with a stretch of Ns. A list of nuclear encoded proteins that physically interact with mitochondrial genes was derived from the *Saccharomyces* Genome Database (SGD, www.yeastgenome.org).

### Functional annotation

Gene ontology analysis was performed using the DAVID functional annotation tools (Huang et al. 2009; Jeong et al 2000) with a fold-change of ≥ 2 at p ≤ 0.05.

### Co-expression analysis

Based on Pearson correlation of the allele expression profiles, a co-expression network was constructed, and clusters of genes/alleles were identified that showed similar expression patterns in the four testing conditions and the two hybrids. The average path length of this network is 9.86, and node degree ranges between 1 and 26 with significant fit to a power law distribution (exponent = 2.43). This indicates that the network has a scale-free property which is common to most biological systems including metabolic pathways and transcriptional regulation maps (Jeong et al 2000; Albert 2005). This also implies that the network has a modular structure in which genes/alleles are co-regulated in response to changes in medium, temperature and mitotype (Thieffry and Romero 1999).

## Acknowledgements

SHK was supported by the Biotechnology and Biological Sciences Research Council DTG programme (BB/F017227/1). KD scholarship was funded by the Royal Government of Thailand under the scheme for Development and Promotion of Science and Technology Talent (DPST) projects. KD was also supported by Wellcome Trust (104981).

